# Effect of glycogen synthase kinase-3 inactivation on mouse mammary gland development and oncogenesis

**DOI:** 10.1101/001321

**Authors:** Joanna Dembowy, Hibret A. Adissu, Jeff C. Liu, Eldad Zacksenhaus, James R. Woodgett

## Abstract

Many components of Wnt/β-catenin signaling pathway have critical functions in mammary gland development and tumor formation, yet the contribution of glycogen synthase kinase-3 (GSK-3α and GSK-3β) to mammopoiesis and oncogenesis is unclear. Here, we report that WAP-Cre-mediated deletion of GSK-3 in the mammary epithelium results in activation of Wnt/β-catenin signaling and induces mammary intraepithelial neoplasia that progresses to squamous transdifferentiation and development of adenosquamous carcinomas at 6 months. To uncover possible β-catenin-independent activities of GSK-3, we generated mammary-specific knock-outs of GSK-3 and β-catenin. Squamous transdifferentiation of the mammary epithelium was largely attenuated, however mammary epithelial cells lost the ability to form mammospheres suggesting perturbation of stem cell properties unrelated to loss of β-catenin alone. At 10 months, adenocarcinomas that developed in glands lacking GSK-3 and β-catenin displayed elevated levels of γ-catenin/plakoglobin as well as activation of the Hedgehog and Notch pathways. Collectively these results establish the two isoforms of GSK-3 as essential integrators of multiple developmental signals that act to maintain normal mammary gland function and suppress tumorigenesis.

## Introduction

Epithelial malignancies, including those of the breast, are thought to initiate in stem-like cells defined by their capacity to self-renew and persist long enough to sustain and propagate mutations, as well as generate all functional cell types within a given tissue. These ‘tumor initiating cells’ (TICs) may arise from normal adult stem cells as a consequence of alterations in the regulation of balance between self-renewal and differentiation or from more committed progenitors that have re-acquired stem cell characteristics during transformation (1, 2). The contribution of distinct stem/progenitor cells to breast cancer heterogeneity remains obscure. With the identification of candidate mammary stem and progenitor populations that vary in their differentiation capacities, the nature of the TIC population and the impact of oncogenes within this critical cellular pool are being elucidated (1, 3). Multiple lines of evidence have shown developmental pathways including Wnt, Hedgehog (Hh) and Notch regulate stem cell homeostasis and have the capacity to induce cancer of various tissues when upregulated. The protein-serine kinases GSK-3α and β are shared components of these networks and act to regulate signals emanating from receptors upon morphogen or ligand ligation. Inhibition of GSK-3 is critical to canonical Wnt signaling leading to increased β-catenin levels associated with elevated proliferation and suppression of differentiation in a number of tissues (4–7). GSK-3 has been shown to interact with Hh pathway components at several levels to inhibit transcriptional functions of Gli proteins (8–11). Several cell line studies have demonstrated that GSK-3 inhibition also modulates Notch signaling, at least in part by phosphorylating the Notch intracellular domain (NICD), thereby preventing nuclear entry and efficient association with its target gene promoters (12–14).

Pathological hyperactivation of each of these pathways is a hallmark of many tumors, including breast cancer. While mutations in β-catenin and other pathway components, such as adenomatous polyposis coli (Apc), are some of the most frequent signaling abnormalities contributing to human tumor pathogenesis, they are much less frequent in breast cancer. Despite the rarity of mutations, increased cytoplasmic and nuclear β-catenin levels have been documented in ∼40% of primary breast cancers including metaplastic breast carcinomas (MBCs), a rare yet aggressive subset of breast cancer that shares features of both basal-like and triple negative breast cancers (TNBCs) (15–20). The mechanism by which Wnt/β-catenin signaling is activated in these tumors is unknown but may involve excess Wnt ligand production and aberrant receptor activation (21–23). Wnt/β-catenin functions as a stem cell survival factor in continuously self-renewing systems including mammary tissues (24, 25). Non-neoplastic glands of MMTV-Wnt1 and those from β-catenin mutant mice where the transgene lacks the N-terminal region containing GSK-3 regulatory sites present an expanded progenitor cell fraction and limited or impaired capacity for functional differentiation in favor of a less committed cellular state (26, 27). In addition, Wnt/β-catenin-responsive cells have recently been shown to be long-lived stem cells that make up a large proportion of the basal compartment able to survive multiple rounds of lobuloalveolar tissue turnover (28). This pool of vulnerable cells may constitute a vulnerable population for oncogenic mutation leading to generation of TICs. Indeed, stabilization of a β-catenin transgene expressed from its endogenous promoter or via Apc mutations or deficiency leads to mammary hyperplasias, accompanied by loss of alveolar structures (29–32), while overexpression of N-terminally truncated β-catenin results not only in precocious alveolar differentiation but also in formation of mammary adenocarcinomas (33–35). In addition, overexpression of the positive Wnt/β-catenin regulator, casein kinase 2 (CK2), as well as a putative Wnt/β-catenin target gene, cyclin D1, in the mammary epithelium results in hyperplasias, squamous differentiation and adenocarcinomas (36, 37).

High levels of Notch-1 were found to be associated with a poorer outcome in breast cancer patients while rearrangements of Notch-1 and 2 were found in TNBCs where the fusion transcripts retained exons that encode the NICD thus maintaining transcriptional outputs (38–40). Increased numbers of breast stem/progenitor cells have been reported upon enhanced Notch signaling (41–43). This expansion of immature cells was predictive of tumor formation suggesting NICD1 expression in the mammary epithelium generates a population of unstable, pre-malignant progenitor cells (43). Notch activity has also been shown to result in luminal progenitor expansion leading to hyperplasia and tumorigenesis (41, 44, 45). Recently two Notch-2 progenitor populations were identified within the luminal compartment representing unique mammary lineages (46). Thus, dysregulation of Notch activity may amplify a distinct stem/progenitor population within the mammary tissues, which may contribute to TIC formation.

Hedgehog (Hh) ligand overexpression is associated with the basal-like breast cancer subtype and poor outcome in terms of metastasis and breast cancer-related death while expression of Hh receptor Ptch1 and transcriptional activator Gli1 has been detected in invasive carcinomas but not in normal breast epithelium (47–51). Hh may increase the proliferative capacity and enhance self-renewal of mammary progenitors rather than enlarge the multi-potent stem cell pool, thus perturbing the stem cell vs progenitor cell balance (52–55). In concordance, Gli1 overexpression within the mammary epithelium induces mammary tumors with increased levels of Bmi-1, a transcriptional repressor of the polycomb gene family previously implicated in stem cell maintenance (56, 57).

To assess the functional consequence of GSK-3 loss from the mammary epithelium and to explore potential β-catenin-independent roles of these protein kinases, both GSK-3 genes were deleted specifically in mammary tissues in the presence or absence of a functional β-catenin gene. Glands of GSK-3 mutant animals assumed epidermal identity via cell-autonomous mechanisms leading to adenosquamous carcinoma formation. Although mammary-selective inactivation of β-catenin initially subverted epidermoid transdifferentiation of GSK-3 null tissue, misregulation of Hedgehog and Notch pathways contributed to the development of adenocarcinomas in the absence of GSK-3 and β-catenin. Thus, GSK-3, by restraining multiple pathways maintains mammary epithelial cell function and inhibits tumor formation.

## Results

### Generation of mice harboring mammary-gland selective deletion of GSK-3α and GSK-3β

To investigate the functions of GSK-3s in the mammary gland, we generated GSK-3α^-/-^; GSK-3β^Exon2^ ^LoxP/Exon2^ ^LoxP^, referred to as GSK-3α^-/-^; GSK-3β^FL/FL^ mice with mammary gland-selective deletion of the floxed GSK-3β alleles achieved by Cre-mediated recombination driven by the whey acidic protein (WAP) promoter yielding WAP-GSK-3 <u>d</u>ouble <u>k</u>nock-<u>o</u>ut animals (WAP-DKO). Expression of the WAP-Cre transgene can be detected during the estrus stage of the murine estrus cycle (hence starting at ∼4 weeks of age) and the WAP promoter is substantially increased during each pregnancy when WAP expression is observed in both ductal and alveolar epithelial cells, particularly starting from 15 days post-coitum (dpc 15) (58–60).

Genomic PCR of WAP-DKO pregnant glands confirmed absence of GSK-3α and loss of GSK-3β (Fig 1A). Cre expression was found in a large proportion of WAP-DKO cells as assessed by IHC (Fig 2). Immunoblotting of pregnant WAP-DKO whole-gland lysates demonstrated excision of the GSK-3β^FL/FL^ allele (Fig 1B). Unrecombined GSK-3β present in the non-epithelial, stromal compartment (e.g. fat, extracellular matrix, immune cells) of mammary tissues not targeted by WAP-Cre likely accounts for the observed residual GSK-3β protein.

**Figure 1:**
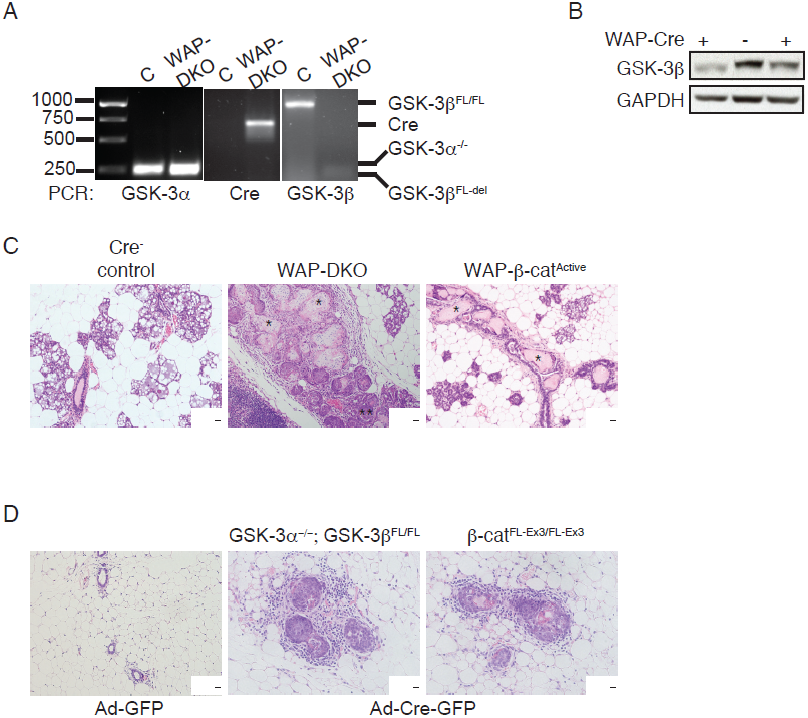
Loss of GSK-3 in the mammary epithelium results in mammary intraepithelial neoplasia (MIN) and squamous transdifferentiation. (**A**) Mid-pregnant glands were isolated from either GSK-3α^-/-^; GSK-3β^FL/FL^ (control, C) or WAP-Cre; GSK-3α^-/-^; GSK-3β^FL/FL^ (WAP-DKO) mice and were PCR-analyzed using primers specific for the detection of floxed (850 bp) and deleted (250 bp) alleles of GSK-3β, Cre (750 bp) or GSK-3α (250 bp) confirming GSK-3α null status and excision at the GSK-3β locus. (**B**) Representative immunoblot of WAP-DKO whole gland lysates probed with indicated antibodies indicating depletion of GSK-3β protein in two Cre positive animals. GAPDH was used as a loading control. (**C**) H&E staining of glands removed from control, WAP-DKO (n=14) or WAP-β-catenin^Active^ (n=6) mice as indicated. The degree of transdifferentiation varied across glands; shown are representative images. *Denotes ghost cells; a region of MIN is denoted by ** in the middle panel. (**D**) Mammary gland outgrowth produced by injection of dissociated control (Ad-GFP), GSK-3 ablated or β-catenin-stabilized (Ad-Cre-GFP) MECs into a cleared fat pad. Squamous foci were observed in 2/3 DKO and 3/3 β-catenin^Active^ MEC-transplanted glands indicating these effects are cell-autonomous and intrinsic to the mammary epithelium.

**Figure 2:**
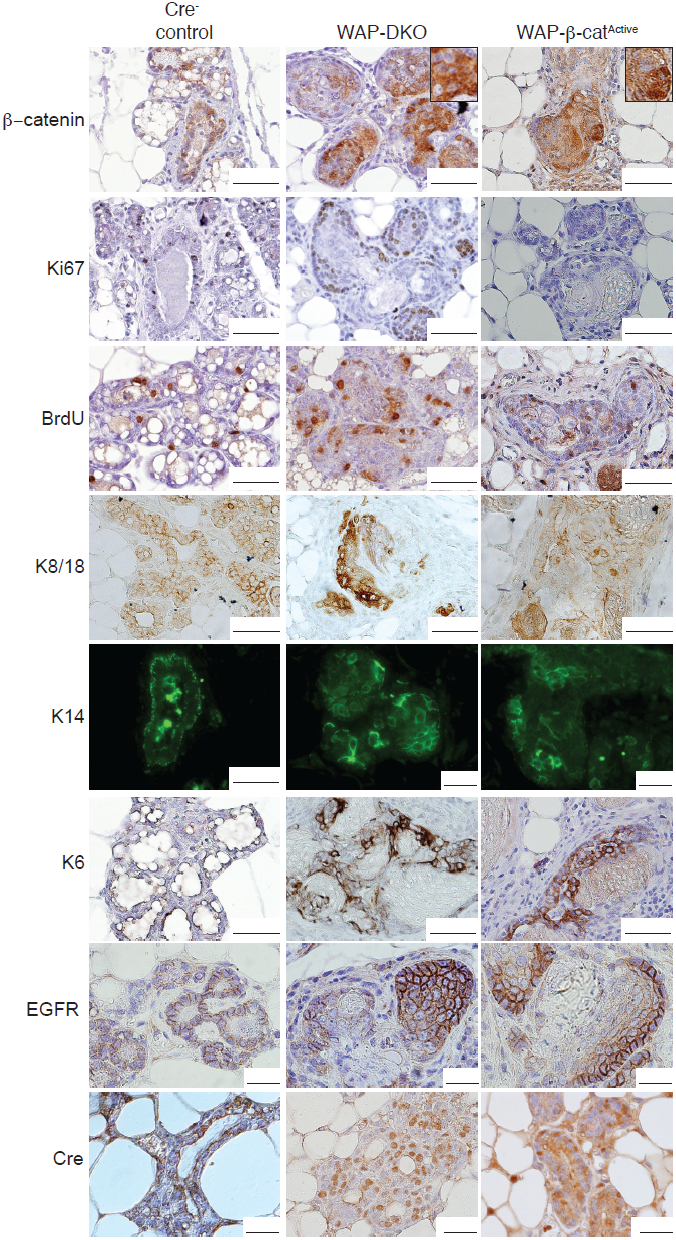
Comparison of mammary glands from WAP-DKO and WAP-β-catenin^Active^ mid-pregnant animals. Representative immunostained sections of glands removed from primiparous animals of indicated genotypes at 15 days post-coitum (dpc 15). Tissues were probed with antibodies to identify cells bearing mammary lineage markers (luminal K8/18, myoepithelial K14), keratin 6 (K6, see text) and to assess proliferation (BrdU, Ki67) and levels of β-catenin, EGFR and Cre. Similar staining patterns across this marker panel were observed in both WAP-DKO and WAP-β-catenin^Active^ pregnant glands with strong nuclear and cytoplasmic β-catenin (insets) and high proliferation as well as presence of both luminal and myoepithelial cells that also strongly express EGFR. Cre expression varied across glands and representative images are shown. Total number of glands analyzed: WAP-DKO, n=5; WAP-β-catenin^Active^, n=3, Cre^-^ controls (includes GSK-3α^-/-^; GSK-3β^FL/FL^ and β-catenin^Active^), n=5.

β-catenin is a major GSK-3 target implicated in multiple stages of mammary development as well as in oncogenesis. For direct comparison with GSK-3 DKO animals, we also generated mice with stabilized β-catenin, which have been characterized previously (31). In this case, endogenous β-catenin was activated in mammary tissues by WAP-Cre recombination of the floxed exon 3 allele of β-catenin (β-catenin^Exon3^ ^LoxP/+^, referred to as β-catenin^FL-Ex3/+^) that encodes for the portion of the amino-terminus containing GSK-3 regulatory phosphodegron sites, thus termed WAP-β-catenin^Active^.

### WAP-Cre-mediated inactivation of GSK-3 in the murine mammary epithelium results in squamous transdifferentiation

Approximately 81% of primi-parous midpregnant (dpc 15) WAP-DKO glands exhibited pronounced epidermoid transdifferentiation (Table 1, Fig 1C), similar to that found in WAP-β-catenin^Active^, in agreement with previous reports (31, 61). The mosaic distribution of WAP-Cre, reflecting the heterogeneity in the synthetic activities of cells within individual alveoli during estrous and at midpregnancy, likely accounts for the variability in the phenotypes observed and also strongly suggests these effects are cell-autonomous. Mammary intraepithelial neoplasia (MIN) was a predominant feature of WAP-DKO glands where lumens of alveoli were filled with atypical cells in a solid pattern. Multilayered structures exhibiting squamous differentiation with central ghost cells (large masses of ‘shadow’ cells or cellular ghosts lacking nuclear and cytoplasmic details with clear conservation of basic cellular outline) and keratinization were also frequently observed and appeared to arise from within MIN lesions (Fig 1C). In contrast, MIN was rare in WAP-β-catenin^Active^ glands, which were found to contain squamous differentiation as well as prominent keratinization with many of the larger lesions completely replaced by ghost cells (Fig 1C). Whereas lesions in dpc 15 WAP-DKO glands appeared to arise from both the alveolar and ductal secretory epithelium, ducts seemed to be preferentially affected in WAP-β-catenin^Active^ (Fig 1C). Unsurprisingly, the transdifferentiated glands of WAP-DKO and WAP-β-catenin^Active^ females were unable to support nourishment of pups and litters were found dead within 24 hr of birth with no evidence of milk spots.

**Table 1.** Spectrum of phenotypes observed in females across examined models.

| Stage | Dpc 15 | Multiparous |  |
| --- | --- | --- | --- |
| Genotype | Incidence of transdifferentiation | Median tumor latency (days) | Metastasis |
| **WAP-DKO** | 75% (n=12) | 180.5 (n=18) | 0 (n=10) |
| **WAP-TKO** | 40% (n=25)* | 300 (n=27) | 0 (n=13) |
| **WAP-β-cat<sup>Active</sup>** | 100% (n=3) | 211 (n=10) | 11% (n=18) |
| **$\frac{3}{4}$ WAP-DKO** | NA | 429 (n=4) | 0 (n=3) |

\*Basal background level of very mild single foci also observed in some Cre<sup>-</sup> controls

To examine the nature of effects of GSK-3 deletion on mammary gland development, we enriched for primary mouse mammary epithelial cells (MECs) from GSK-3α^-/-^; GSK-3β^FL/FL^ mice and infected them with adenovirus expressing Cre-GFP (Ad-Cre-GFP) or GFP alone (Ad-GFP) as a control. We then introduced paired groups of KO and control MECs into the contralateral cleared mammary fat pads of 3-wk-old FVB mice and allowed them to develop for a period of 8 wks. After infection with Ad-GFP, floxed MECs generated normal mammary epithelial outgrowths upon transplantation (Fig 1D). In contrast, after infection with Ad-Cre-GFP, squamous foci were once again observed in fat pads transplanted with GSK-3 KO MECs (Fig 1D). Similarly, epidermoid transdifferentiation was also found in outgrowths from β-catenin^Active^ MECs following injection of Ad-Cre-GFP-infected β-catenin^FL-Ex3/+^ MECs (Fig 1D). These data provide evidence that effects of mammary tissue-specific ablation of GSK-3 are intrinsic to the mammary epithelium.

### Activation of Wnt/β-catenin signaling in mammary glands lacking GSK-3

More detailed analysis revealed a correlation between the level of β-catenin expression and changes in cell morphology in WAP-DKOs and in WAP-β-catenin^Active^ (Fig 2), as β-catenin was primarily found surrounding keratinized structures. Nuclear β-catenin was never observed in control (GSK-3α^-/-^; GSK-3β^FL/FL^) virgin glands in which it is normally maintained at a low level and is localized primarily at the membrane (Fig 2). As assessed by increased BrdU incorporation and Ki67 staining, dpc 15 WAP-DKO glands were actively proliferating (Fig 2), beyond the level associated with normal mammary growth during pregnancy. Variable expression of cytokeratin 8/18 (K8/18) and cytokeratin 14, markers of luminal and myoepithelial lineages respectively, was maintained within the MIN lesions (Fig 2). Keratinizing epithelium of both WAP-DKO and WAP-β-catenin^Active^ was associated with intense expression of cytokeratin 6 (K6) (Fig 2), which was never observed in control glands. Based on its expression pattern, which is observed early in embryonic mammary development and in non-proliferating terminal end buds but rare in mature glands, K6 is considered a putative progenitor marker in the mammary epithelium (27, 62, 63). However, since K6 is also found in activated keratinocytes in the epidermis (63), it is difficult to discern whether the observed elevation of K6 in WAP-DKO and WAP-β-catenin^Active^ represents an expansion of immature cells or is a consequence of epidermoid transdifferentiation. Flow cytometric analysis of MECs purified from dpc 15 glands of both strains using CD24 and CD49f mammary stem cell markers did not reveal abnormalities in the stem cell profile where the ratios of luminal (CD24^hi^:CD49f^+^) to basal (CD24^lo^:CD49f^hi^) cells were similar to control glands (data not shown). This result is consistent with positive K6 staining reflecting a switch to epidermoid marker expression in transdifferentiated cells rather than aberrations within the mammary stem cell compartment. These observations correlate β-catenin stabilization with proliferation, neoplastic lesion formation and alterations in the differentiation status of the mammary epithelium. Taken together these results show that the two GSK-3 isoforms, by restraining β-catenin levels and ensuing proliferation, play a key role in maintaining glandular cell fate in the mammary epithelium.

### Loss of GSK-3 and activation of β-catenin have differential effects on mammary stem cells

To further investigate GSK-3 functions we employed a mammosphere (MS) assay to measure *in vitro* stem/progenitor cell frequency in primary MEC preparations (64). In the absence of attachment to an exogenous substratum or cell-cell adhesion, stem cells are able to survive and proliferate and we found no significant differences in primary or secondary MS formation of GSK-3 DKO MECs generated by infecting Lin^-^ GSK-3α^-/-^; GSK-3β^FL/FL^ cells with either Ad-GFP or Ad-Cre-GFP. Interestingly, whereas the CD24:CD49f marker profile of MS dissociated on day 12 post-infection was not significantly changed upon loss of GSK-3 across replicate experiments (Fig 3B,C) a prominent CD24^hi^:CD49f^lo/-^ population was observed upon stabilization of β-catenin (Fig 3A,C). To further investigate the functional relevance of these populations, β-catenin^Active^ MECs were sorted into CD24^hi^:CD49f^lo/-^ and CD24^lo/-^:CD49f^hi^ fractions with only CD24^lo/-^:CD49f^hi^ cells capable of forming secondary MS, although CD24^hi^:CD49f^lo/-^ MEC population was regenerated (data not shown). Based on these *in vitro* data, we conclude that, unlike direct stabilization of β-catenin, GSK-3 loss does not significantly impact the mammary gland stem cell compartment.

**Figure 3:**
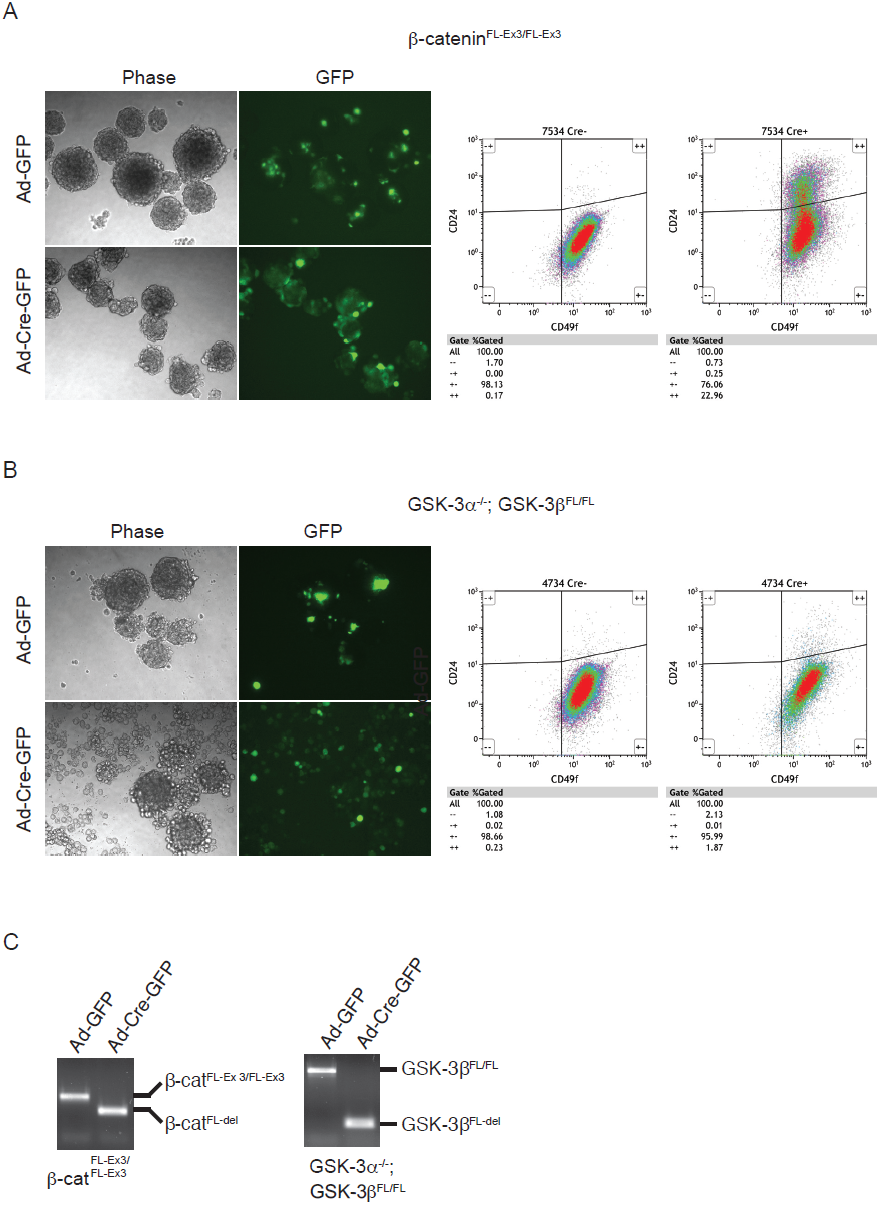
Mammosphere formation in MECs depleted of GSK-3 and β-catenin. β-catenin^Active^ **(A)** or GSK-3-ablated **(B)** MECs were cultured under conditions promoting MS formation and imaged on day 12 post-infection. MS were then dissociated and analyzed by flow cytometry to assess the CD24:CD49f stem cell profile. Shown are representative of 3 separate experiments demonstrating efficient MS formation across all genotypes. A CD24^hi^:CD49f^lo/-^ population was consistently observed in β-catenin^Active^ MECs. **(C)** PCR-genotyping of an aliquot of Ad-GFP or Ad-Cre-GFP MECs of indicated genotypes analyzed using primers specific for detection of floxed and deleted alleles of GSK-3β (850 bp, 250 bp) and β-catenin^FL-Ex3/FL-Ex3^ (650 bp, 400 bp) showing efficient Cre excision at both loci at day 12 post-infection.

### Mammary tumors develop in absence of GSK-3

Following a median 6-7 months of continuous breeding, all WAP-DKO and WAP-β-catenin^Active^, but not control female mice, developed large-sized mammary ductal adenocarcinomas with atypical squamous transdifferentiation (Table 1, Fig 4C). There was no statistical difference in survival curves for these strains (P=0.2333 log-rank test, P=0.3706 Wilcoxon test) (Fig 4C). Near complete loss of GSK-3β in WAP-DKO tumors was confirmed by detection of markedly elevated β-catenin levels by immunoblotting (Fig 4A).

**Figure 4:**
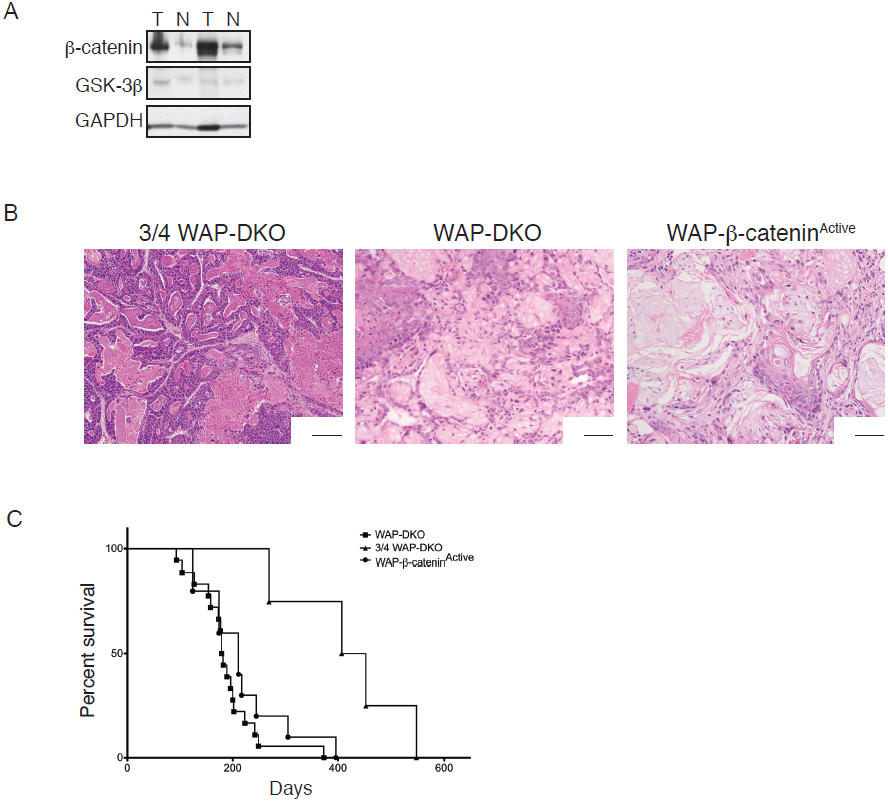
Multiparous WAP-DKOs and WAP-β-catenin^Active^ develop adenosquamous carcinomas. **(A)** Immunoblots of whole-tissue lysates of tumors harvested from 2 separate multiparous WAP-DKO animals showing loss of GSK-3β and significant elevation of β-catenin specifically in the tumors (T) when compared with adjacent normal mammary tissue (N). Total number of tumor-bearing animals analyzed, n=8. **(B)** H&E staining of tumors arising in transgenics of indicated genotypes showing corresponding histopathology. Total number of tumors analyzed: WAP-DKO, n=18; WAP-β-catenin^Active^, n=10. **(C)** Kaplan-Meier survival analysis of tumor-bearing transgenic animals used in the study showing separation of WAP-DKO (n=18) and WAP-β-catenin^Active^ (n=10) cohorts from ¾ WAP-DKOs (n=4). Age of death represents the moment at which, due to the presence of signs of discomfort or because the tumor size exceeded 2 cm^3, mice had to be euthanized according to institutional and national regulations.

The primary tumors in WAP-DKO and WAP-β-catenin^Active^ were composed of poorly differentiated glandular structures of variable size and large numbers of multilayered squamous epithelial structures (Fig 4B). The neoplastic cells had a high nuclear-to-cytoplasmic ratio (1:1), a moderate amount of granular basophilic cytoplasm and a round to oblong large nucleus with fine chromatin and 1-3 prominent nucleoli. There was marked anisocytosis (variation in cellular size) and anisokaryosis (variation in nuclear size). Occasional, large multinucleated cells were also present. Nearly half of the tumor volume was composed of multilayered epithelial structures with squamous differentiation and central ghost cells. Some of these structures contained lamellar keratin material and in rare areas had features reminiscent of normal stratified, keratinized, squamous epithelium (epidermis). Invasion to the regional lymph node was observed occasionally in WAP-DKO and WAP-β-catenin^Active^ tumors (data not shown).

As expected, β-catenin was elevated specifically in WAP-DKO tumors as compared to adjacent normal mammary tissue (Fig 4A). Tumor expression of ducto-luminal (K8/18) and ducto-basal (K14) cells was maintained along with K6 and EGFR (Fig 5). A high level of proliferation was evident in all tumors based on Ki67 staining (Fig 5).

**Figure 5:**
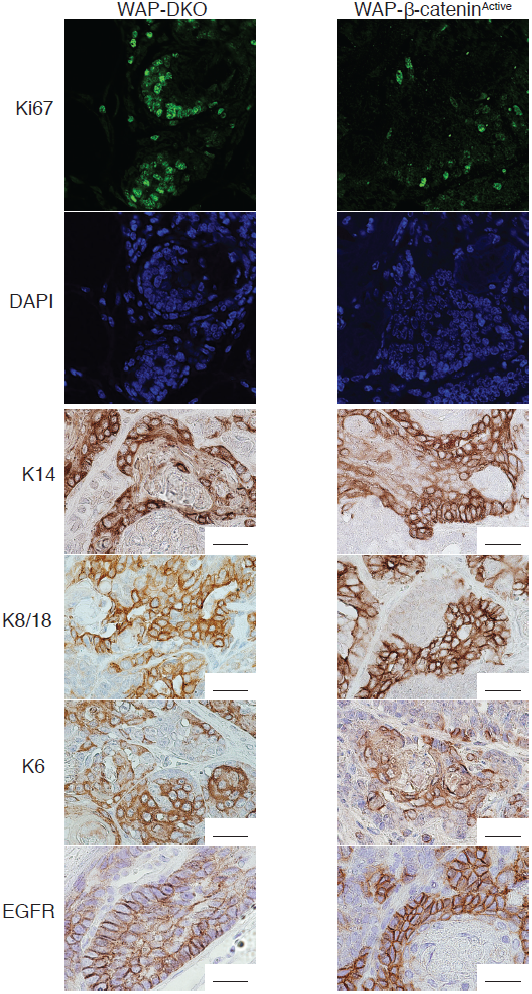
WAP-DKO and WAP-β-catenin^Active^ tumors display Wnt pathway activation, express luminal and basal markers and EGFR. Representative immunostained sections of tumors harvested from animals of indicated genotypes showing nuclear and cytoplasmic β-catenin indicative of Wnt pathway activity. Based on marker expression, tumors from both WAP-DKOs and WAP-β-catenin^Active^ transgenics are EGFR positive and contain luminal and myoepithelial/basal cells. Number of tumors analyzed: WAP-DKO, n=6 and WAP-β-catenin^Active^ (n=3).

We found ∼10% rate of pulmonary metastasis in WAP-β-catenin^Active^ while none of the WAP-DKO tumors were ever found to metastasize (Table 1). Further assessment of metastatic potential showed Lin^-^ tumor cells from 2 of 8 WAP-β-catenin^Active^ were able to regenerate tumors when injected into syngeneic recipients suggesting their weak malignant propensity (data not shown). In contrast, none of WAP-DKO Lin^-^ tumor cells grew following transplantation.

Tumor incidence was also noted in 20% of glands lacking 3 of the 4 GSK-3 genes, i.e. of genotype WAP-Cre; GSK-3α^-/-^; GSK-3β^FL/+^; or WAP-Cre; GSK-3α^-/+^; GSK-3β^FL/FL^ (referred to as ¾ WAP-DKO) arising with a median latency of 14 months, which was statistically significantly different from WAP-DKO (P=0.0015 log-rank test, P=0.0075 Wilcoxon test) (Table 1, Fig 4C). No tumors were observed in animals with either just the α or β isoform of GSK-3 missing. The markedly different morphology of ¾ WAP-DKO tumors (Fig 4B) and their low incidence may reflect these being spontaneous age-related tumors previously reported in the FVB strain rather than attributable to loss of GSK-3. This may also be true of the single observed case of tumor formation in virgin WAP-DKOs (latency ∼13.6 months). However, we cannot exclude the possibility that loss of 3 of the 4 alleles causes weak tumor predisposition.

From these data, we conclude that depletion of all four alleles of GSK-3 from the mouse mammary epithelium leads to rapid formation of adenosquamous carcinomas indicating GSK-3 functions to inhibit tumorigenesis within this compartment.

### Loss of GSK-3 drives β-catenin-independent mammary tumorigenesis

To distinguish possible β-catenin-independent GSK-3-specific functions, mice harboring LoxP sites in introns 2 and 6 of β-catenin (β-catenin^Exon^ ^2-6^ ^LoxP/Exon^ ^2-6^ ^LoxP^, referred to as β-catenin^FL-Ex2-6/FL-Ex2-6^) were mated with WAP-DKOs to generate animals with mammary glands lacking both pairs of GSK-3 alleles as well as β-catenin (referred to as WAP-triple knockout, WAP-TKO) (Fig 6A,B). Residual unrecombined alleles of GSK-3β and β-catenin in mid-pregnant WAP-TKOs mammary compartments lacking expression of WAP-Cre likely reflect presence of these proteins in whole gland lysates (Fig 6B). Strikingly, dpc 15 WAP-DKO squamous transdifferentiation was rescued by loss of β-catenin and no alterations in stem cell populations were observed as judged by the CD24:CD49f profile (data not shown). To determine whether simultaneous ablation of GSK-3 and β-catenin affected the propensity of MECs to form mammospheres, GSK-3α^-/-^; GSK-3β^FL/FL^; β-catenin^FL-Ex2-6/FL-Ex2-6^ MECs were infected with either Ad-Cre-GFP or Ad-GFP. Whereas MS formation was unaffected by Ad-GFP, no spheres were ever observed in TKO MECs (Fig 6C). Analysis of the CD24:CD49f stem cell profile of these TKO MECs did not reveal any aberrations (Fig 6C). To evaluate the possibility that membrane/adhesion functions of β-catenin are necessary for sphere formation, we subjected β-catenin^FL-Ex2-6/FL-Ex2-6^ MECs to the MS assay and found sphere-forming ability was normal (data not shown). Thus, in this short-term assay, loss of GSK-3 and β-catenin abrogates stem cell functionality without perturbation to the stem cell compartment.

**Figure 6:**
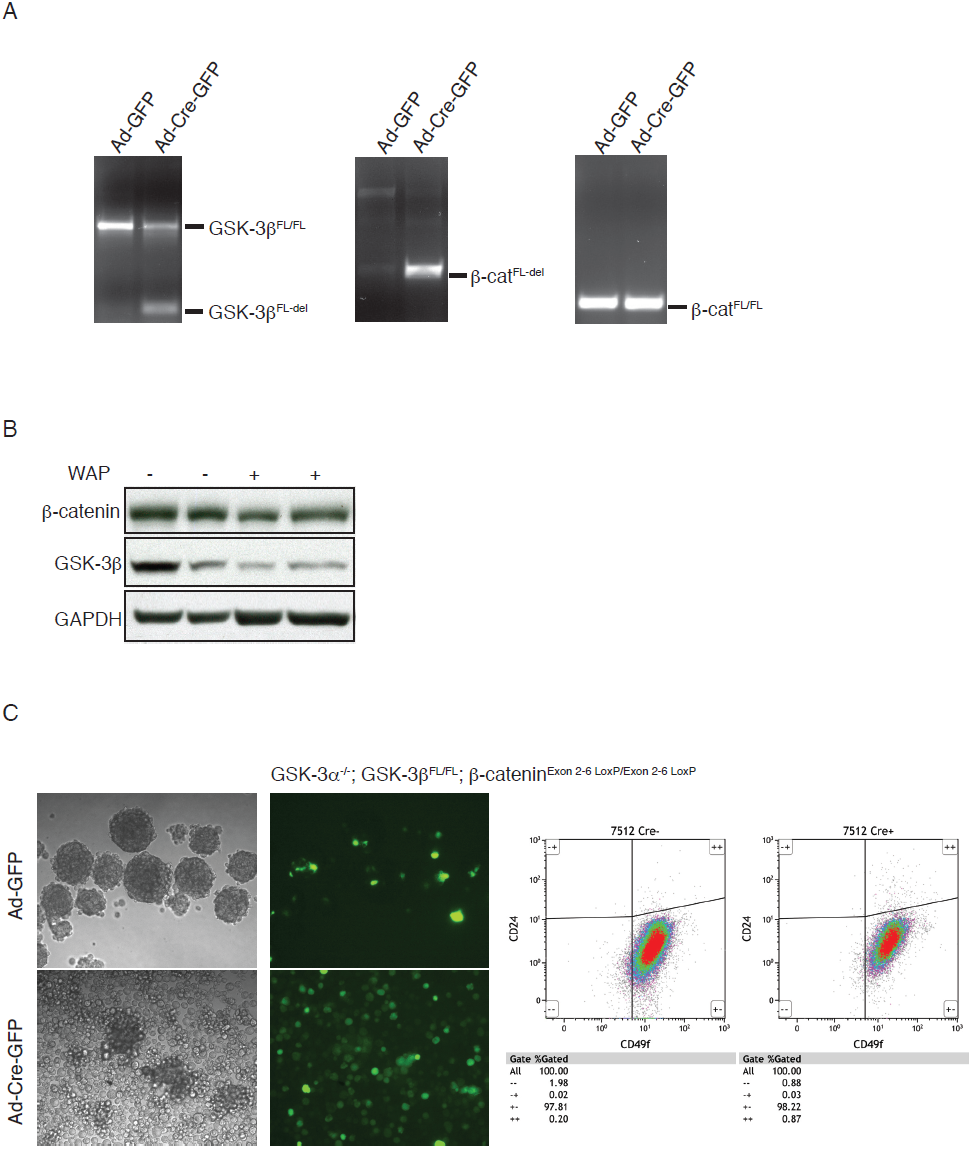
Mammary-selective deletion of GSK-3 and β-catenin (WAP-TKO) rescues epidermoid transdifferentiation and inhibits mammosphere formation. **(A)** PCR-based assessment of floxed and excised alleles of GSK-3β (850 bp, 250 bp) and β-catenin^FL-Ex2-6/FL-Ex2-6^ (324 bp, 631 bp) in mid-pregnant mammary glands from GSK-3α^-/-^; GSK-3β^FL/FL^; β-catenin^FL-Ex2-6/FL-Ex2-6^ mice mediated by WAP-Cre. **(B)** Representative immunoblot of whole gland lysates from dpc 15 WAP-TKO mice probed with indicated antibodies showing reduced GSK-3β and β-catenin protein levels. **(C)** Whereas MS formation proceeded normally in Ad-GFP-infected GSK-3α^-/-^; GSK-3β^FL/FL^; β-catenin^FL-Ex2-6/FL-Ex2-6^ MECs, no spheres formed when β-catenin and GSK-3 were deleted simultaneously. The CD24:CD49f stem cell profile was not affected in TKO MECs. Shown are representative of 3 separate experiments.

After ∼10 months, multiparous WAP-TKO dams developed mammary tumors (Table 1, Fig 7A). This temporal incidence was statistically significantly different from both WAP-DKO and WAP-β-catenin^Active^ tumor latency (P<0.0001 log-rank and Wilcoxon tests) (Fig 7A). Unlike WAP-DKOs, which developed adenosquamous carcinomas with a prominent squamous component, acinar adenocarcinomas formed in WAP-TKOs with rare and multifocal squamous foci present in some tumors (Fig 7C). Both mammary lineage markers continued to be expressed in these tumors along with K6, SMA, EGFR and Ki67 (Fig 7E). GSK-3β expression was virtually undetectable in these tumors (Fig 7B) and laser-capture micro-dissection (LCM) demonstrated efficient excision at the β-catenin locus in both the adeno and squamous components of tumors (data not shown). Interestingly we found elevated levels of γ-catenin/plakoglobin (Fig 7B), an armadillo domain-containing β-catenin homologue that associates with both classic and desmosomal cadherins (65) specifically in WAP-TKO tumors suggesting it may compensate for loss of β-catenin functions, at least at the level of cell:cell adhesion. In addition to its role in the Wnt/β-catenin pathway, GSK-3 also coordinates proliferation and differentiation signals through regulation of critical nodes in PI3K, Hh and Notch signaling pathways. While we found no change in phosphorylation of PKB/Akt (as a marker for PI3K activation, data not shown), examination of overall levels of Hh effector Gli1 via immunoblotting demonstrated Gli1 to be elevated specifically in WAP-TKO tumors (Fig 7B). To assess the contribution of Notch signaling to WAP-TKO tumor-initiating cell (TIC) activity, Lin^-^ TICs were allowed to form spheres and subsequently treated with escalating doses of γ-secretase inhibitor (GSI) (Fig 7D). Even at low doses of GSI, a reduction in the number of TS was observed while secondary TS formation was completely abrogated (Fig 7D). Taken together, these findings provide evidence that in the absence of GSK-3, strong selective pressure exists for increases in γ-catenin along with induction of major developmental signaling pathways, Hh and Notch, which may underlie β-catenin-independent tumorigenic effects of GSK-3 disruption.

**Figure 7:**
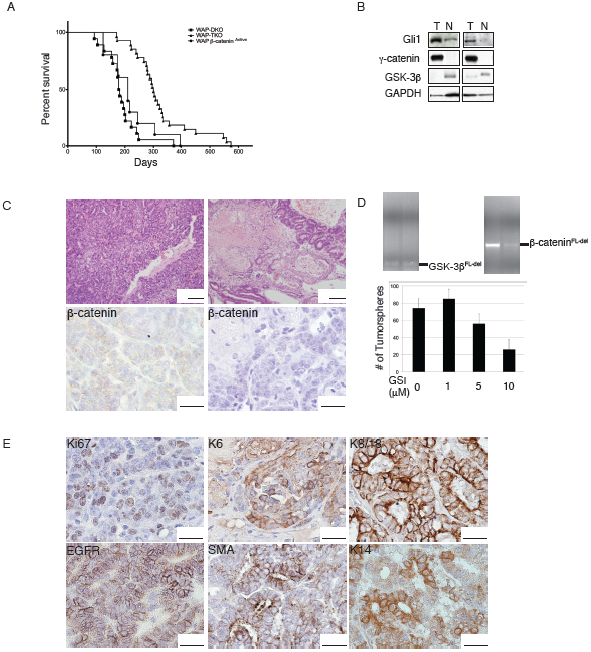
Mixed histopathology of tumors arise following long latency in WAP-TKO mice. **(A)** Kaplan-Meier survival analysis of tumor-bearing transgenics showing separation of WAP-TKOs (n=27) from both WAP-DKO (n=18) and WAP-β-catenin^Active^ (n=10) cohorts. **(B)** Immunoblot showing loss of GSK-3β and elevated levels of Gli1 and γ-catenin specifically in WAP-TKO tumors. GAPDH serves as loading control (T, tumor; N, normal). Total number of tumors analyzed, n=10. **(C)** WAP-TKO tumors are largely adenocarcinomas that contain elements of squamous differentiation (top panels) with no β-catenin staining in either histological component. Total number of tumors analyzed n=27. **(D)** WAP-TKO tumor cells were purified, allowed to form tumorspheres (TS) and subsequently treated in triplicates with increasing concentrations of γ-secretase inhibitor (GSI) as indicated. TS were counted on day 7 of treatment (bottom panel). Efficient excision at the GSK-3β and β-catenin was confirmed by PCR genotyping of TS (top panel) from 2 separate tumors. **(E)** Representative immunostained sections of WAP-TKO tumors showing expression of luminal and myoepithelial/basal markers, proliferation as well as presence of K6, EGFR and SMA. Total number of tumors analyzed, n=8.

## Discussion

GSK-3 is a primary negative regulator of β-catenin and we anticipated similar phenotypes of GSK-3 inactivation compared to genetic activation of β-catenin. Given extensive literature demonstrating that activation of Wnt signaling in mammary epithelium induces squamous differentiation (31, 36, 37, 61, 66), it was not surprising that loss of GSK-3 also conferred squamous potential on mammary cells. This phenotype is cell-autonomous and although strikingly similar to that observed upon direct stabilization of β-catenin, at the stem cell level GSK-3 also exhibits distinct functions. Elevation of β-catenin results in the generation of a CD24^hi^ population not found in the absence of GSK-3 and this may contribute to the malignant character of β-catenin tumors. We speculate that MEC response is largely determined by the specific level of β-catenin and while a lower threshold may be required for epidermoid transdifferentiation common to both WAP-DKOs and WAP-β-catenin^Active^ glands, a stronger β-catenin signal is required for changes associated with malignancy.

### GSK-3 inhibits mammary tumorigenesis by restraining signaling via multiple pathways

Deletion of GSK-3 and the ensuing elevation in β-catenin levels resulted in adenosquamous carcinomas in WAP-DKOs. Although not previously reported, stabilization of endogenous β-catenin in the mammary epithelium also led to tumors in the present study. These WAP-β-catenin^Active^ tumors were histologically similar to those of WAP-DKOs and developed with similar kinetics suggesting that stabilization of β-catenin is the key requisite step for WAP-DKO tumor formation. Thus, elevation of β-catenin may permit MECs to re-enter the cell cycle resulting in MIN lesions that progress to adenosquamous carcinomas.

Analysis of β-catenin-independent tumors formed in absence of GSK-3 revealed markedly elevated expression of γ-catenin/plakoglobin, suggesting a potential compensation mechanism for loss of β-catenin signaling and/or for its role in cell adhesion. This is consistent with recent findings in embryonic stem cells where γ-catenin was found to be responsive to GSK-3 inhibition and can activate TCF target genes when the levels of its expression reach a critical threshold (67). However, despite their interactions with common cellular partners, β-catenin has well-documented oncogenic potential while γ-catenin was characterized as having tumor suppressor activity that may be independent of its role in mediating cell-cell adhesion (68–70). Human tumor analysis has shown loss or reduced expression of γ-catenin is associated with poor clinical outcome and increased tumor formation and metastasis (71–74). The capacity of γ-catenin to suppress motility and migration may also underlie the lack of metastasis observed in WAP-TKOs and possibly the observed lack of MS formation in TKO MECs.

Despite what may be discerned from the volume of literature linking GSK-3 and Wnt, this protein kinase acts as a nexus in the control of several critical cellular homeostatic mechanisms, in addition to those supplied by Wnt/β-catenin. Our results suggest that GSK-3 plays an important role in mediation of Hh signaling as loss of GSK-3 resulted in elevated Gli1 expression and associated signaling. In addition, enhanced Notch signaling in the absence of GSK-3 may contribute to tumor formation through dysregulation of TIC control. We anticipate that effects of GSK-3 on Wnt, Notch and Hedgehog signaling are ongoing however different proportions of the total GSK-3 pool may participate in the regulation of these pathways. GSK-3 is known to localize to different cellular compartments and may be sequestered in multi-vesicular bodies (MVB) as recently proposed (75). Hence, distribution of GSK-3 may be dependent on cellular context and the events integrated by GSK-3 and the relative strength of their output will determine observed cellular responses. Whereas WAP-TKO tumors showed significant activation of Hh and γ-catenin/plakoglobin, these pathways were essentially unchanged in WAP-DKO tumors suggesting GSK-3 predominantly functions within the β-catenin destruction complex to regulate the Wnt pathway when β-catenin is intact.

## Materials & Methods

### Generation of mouse strains used in the study

All animals were housed at the Toronto Centre for Phenogenomics (www.phenogenomics.ca). GSK-3α^-/-^ and GSK-3β^Exon2^ ^LoxP/Exon2^ ^LoxP^ (referred to as GSK-3β^FL/FL^) mutants and their wild-type (WT) littermates, were generated as previously described (76, 77). GSK-3α^-/-^ and GSK-3β^FL/FL^ mice were 6 generations backcrossed to FVB mice. At weaning, all littermates of mixed genotypes were housed by gender in groups of 3 to 5 per cage under a 12-hour light/dark cycle (lights on at 07.00) with *ad libitum* food (Purina mouse chow) and water. To generate mammary gland-specific GSK-3 knock-out (KO) mice, GSK-3β^FL/FL^ animals were crossed with B6.Cg-Tg(Wap-Cre)11738Mam (obtained from Sean Egan, Hospital for Sick Children), which carries Cre recombinase under the control of the whey acidic protein (WAP) promoter. WAP-Cre; GSK-3β^FL/FL^ animals were then mated with GSK-3α^-/-^ to generate WAP-Cre; GSK-3α^-/-^; GSK-3β^FL/FL^, designated WAP-DKO, denoting them as mammary specific GSK-3 <u>d</u>ouble <u>k</u>nock<u>o</u>ut. WAP-Cre and WAP-DKOs were also crossed to B6.129-Ctnnb1^tm2Kem^/KnwJ (backcrossed onto FVB for 6 generations) bearing the exon 2-6 floxed allele of β-catenin (β-catenin^Exon^ ^2-6^ ^LoxP/+^, referred to as β-catenin^FL-Ex2-6/+^). This enabled generation of WAP-Cre; β-catenin^FL-Ex2-6/FL-Ex2-6^ and WAP-Cre; GSK-3α^-/-^; GSK-3β^FL/FL^; β-catenin^FL-Ex2-6/FL-Ex2-6^, or <u>t</u>riple <u>KO</u> referred as WAP-TKO. WAP-Cre mice were also mated to β-catenin^Exon^ ^3^ ^LoxP/+^ (referred to as β-catenin^FL-Ex3/+^) transgenics (obtained with permission from Makoto M. Taketo and Derek van der Kooy, University of Toronto) previously described (78) to generate WAP-β-catenin^Active^ mice.

### BrdU injection

Intraperitoneal injections of mice were performed using BrdU (BD Pharmingen) at 100 µg/g body weight. Incorporation of BrdU was detected 4 hrs post injection using immunohistochemistry (IHC) as described in ‘Histology and immunostaining of tissue sections’.

### Mammary epithelial single-cell preparation, enrichment and flow cytometry analysis

Mammary tumors or glands from mice sacrificed at indicated developmental time points were dissected and digested in 1X Collagenase/Hyaluronidase (StemCell Technologies 07912) diluted in DMEM-F12 and incubated at 4°C overnight (O/N) and then at 37°C for 30 minutes, passed through a 40 µm cell strainer (BD Falcon) and pelleted by centrifugation at 1000 rpm for 5 min at room temperature (RT). Enrichment of Lin^-^ mammary epithelial cells (MEC) was achieved through selective depletion of hematopoietic, endothelial, and stromal cells using an EasySep kit (StemCell Technologies 19757) according to manufacturer’s directions. Lin^-^ cells were then stained with anti-CD49f clone GoH3 conjugated with either R-phycoerythrin, FITC or APC (CD49f-PE, CD49f-FITC, CD49f-APC), anti-CD24 clone M1/69 conjugated with PE, FITC or PerCP-eFluor 710 (CD24-PE, CD24-FITC, CD24-PerCP-eFluor 710) as well as 7-aminoactinomycin D (7-AAD), SYTOX Blue (Life Technologies) or DAPI for 30 min at RT. HBSS containing 2% FBS (referred to as HF) was used as buffer. Live single cells (fixed FSC-A/FSC-W ratio; 7-AAD, SYTOX or DAPI-negative) were gated for analysis. Flow cytometry acquisition was performed using FACSCalibur (Becton Dickinson) or Gallios (Beckman Coulter) flow cytometer using a 10-colour, 4-laser (488 nm blue, 561 nm yellow [co-linear], 638 nm red, 404 nm violet) at standard filter configuration. Fluorescence-activated cell sorting (FACS) was carried out using FACSAria-13 color Cell Sorter (Becton Dickinson) with 488 nm blue laser at 30 p.s.i. All antibodies were titrated to determine the optimal antibody concentration. Briefly, TICs, MECs or MECs expressing GFP were stained with CD24 and CD49f (conjugated to appropriate fluorophore) and SYTOX Blue or 7-AAD and fluorescence-minus-one controls were used to determine the CD24 and CD49f gates. Flow cytometry analysis was performed using the Kaluza analysis software package (Beckman Coulter).

### Mammosphere (MS) and tumorsphere (TS) culturing *in vitro*

Immediately following EasySep Magnet incubation (see above), a single-cell suspension of Lin^-^ MECs was infected with adenovirus-GFP (Ad-GFP) or adenovirus-Cre-GFP (Ad-Cre-GFP) at a multiplicity of infection (MOI) of 50. Virus was allowed to adsorb to cells for 1 hour at 37°C with gentle agitation every 10 min. Infected MECs (for MS) or enriched TICs (for TC) were then plated on ultra–low attachment plates (Corning, Costar) in DMEM/F-12 HAM medium (Sigma) containing 20 ng/mL basic fibroblast growth factor (hFGF; StemCell Technologies), 20 ng/mL epidermal growth factor (EGF; StemCell Technologies) and B-27 supplement (1:50 dilution, Life Technologies) and cultured in a standard tissue culture incubator. Spheres were dissociated at 7-day intervals as follows: spheres were collected in a single tube, spun down and supernatant removed. MS were resuspended in 100 µL of 0.25% trypsin and incubated for 1 min at RT followed by 1 min of mechanical disruption by passage through a 1 mL syringe with a 26-gauge needle. 900 µL of HF was added to neutralize the trypsin and cells were counted and re-plated. To enumerate spheres, prior to enzymatic and mechanical dissociation, spheres were resuspended in a known volume and a 10^th^ of suspension was transferred to a well of a 96-well plate and all spheres were counted. Images were captured using Hamamatsu camera mounted on a Leica inverted DMIRB microscope equipped with Volocity software.

### Transplantation

Sorted tumor cells or Lin^-^ MECs 48 hr post-Ad-infection were resuspended in DMEM-F12 medium and mixed at a 1:1 ratio with Matrigel (BD Bioscience) on ice. 1 µl of Trypan Blue solution (Life Technologies) was added to all samples to enable visualization during injection. A mixture of Matrigel and 1, 2 or 10×10^3 tumor cells was injected using a Hamilton syringe near the upper #4 mammary fat pads (mfp) of 6-8 week-old virgin FVB mice anesthetized with isoflurane. Similarly, 50×10^^^3 of adenovirus-infected MECs were mixed 1:1 with Matrigel and injected into cleared #4 mfp and analyzed 8 weeks later.

### Histology and immunostaining of tissue sections

Mammary glands (thoracic) were harvested at indicated developmental time points. Tumors were harvested at tumor endpoint. Tissues were fixed in 4% paraformaldehyde O/N at 4°C and placed in 70% ethanol until they were paraffin-embedded. 5 µm paraffin sections were stained with hematoxylin and eosin (H&E). Tissue sections were deparaffinized in two 5 min changes of xylenes, rehydrated in graduated ethanols, and then washed in PBS. With the exception of EGFR staining where 1 mM EDTA (pH 8) was used, antigen retrieval was performed using 10 mM citrate buffer (pH 6) in a microwave at 10 min boiling following 10 min sub-boiling. Slides were cooled for 30 min. For IHC, endogenous peroxidase activity was quenched by incubating sections in 0.3% hydrogen peroxide for 30 min followed by blocking in 5% goat serum (GS) in PBS for 30 min or 5% GS in TBS containing 0.1% Tween-20 (TBST) for EGFR staining. For immunofluorescence (IF), sections were blocked for 30 min in IF buffer (PBS+0.2% Triton X-100, 0.05% Tween-20) containing 2% BSA. Sections were then incubated in primary antibody (Supplementary Table 1) O/N at 4°C in 2% BSA-PBS. Following washes with PBS for IHC or IF buffer for IF, sections were incubated for 1 h at RT with biotinylated (Dako) or Alexa Fluor (Life Technologies)-conjugated secondary antibodies for IHC or IF, respectively. For IHC the peroxidase reaction was carried out using DAB substrate (Dako) and sections were counterstained with hematoxylin and mounted in Faramount (Dako) aqueous mounting medium. For IF, sections were mounted in ProLong Gold Antifade Reagent with DAPI (Life Technologies). IHC images were digitally captured with an Olympus digital DP 12 megapixel color camera mounted on the Olympus BX61 motorized microscope. IF images were captured using a Deltavision deconvolution microscope (Applied Precision Inc. Issaquah, WA), that consisted of an Olympus Inverted IX70 microscope and motorized stage. Images were projected onto a Coolsnap CCD camera and processed through Softworx software that contained iterative deconvolution algorithms.

### Immunoblotting

50 mg of tissue or tumor was lysed in 1 mL of RIPA lysis buffer supplemented with complete protease inhibitor tablet (Roche) and phosphatase inhibitor cocktail (Sigma) using TissueLyser. Lysates were cleared by centrifugation for 15 min at 4°C and protein concentration determined by Lowry assay (Bio-Rad). 10–30 µg of protein lysates containing SDS sample buffer were boiled for 5 min and loaded onto an 8% or 10% SDS–PAGE gel, followed by semi-dry transfer onto polyvinylidene difluoride (PVDF) membrane (Millipore). Blocking was performed for 1 hr at RT in 5% non-fat milk in TBST and membranes were probed with primary antibodies (Supplementary Table 1) O/N at 4°C. Following washes with TBST, membranes were incubated with appropriate HRP-conjugated secondary antibodies (Bio-Rad) for 45 min at RT, washed and exposed to film (Kodak) using ECL reagent (Pierce).

### Genotyping

Mouse tail tips were digested in 200 mM Tris HCl pH 8.4, 500 mM KCl containing NP40, Tween-20 and Proteinase K at 60°C for 2 hr. Genotyping of the mice was performed using one pair of appropriate primers (Supplementary Table 2).

## Acknowledgements

We thank Tao Deng, Hartland W. Jackson and Purna A. Joshi for assistance with mammary gland manipulation and Katrina MacAulay, Brad Doble and Satish Patel for their generation of mice harbouring floxed GSK-3 alleles. The authors wish to thank Tara Paton of The Centre for Applied Genomics (TCAG), The Hospital for Sick Children, Toronto, for assistance with Sanger sequencing and Yanchun Wang of the Centre for Modeling Human Disease Pathology (CMHD) and Cryopreservation Core at the Toronto Centre for Phenogenomics for laser-capture microdissection. This work was supported by grants to JRW from the CBCRA, CIHR (MOP74711) and Terry Fox Foundation team grant to EZ. JD was partially supported by a CBCF (Ontario) doctoral fellowship.

## Author Contributions

JD and JCL designed, performed and analyzed the experiments; HAA performed pathological examination and classification of tumor samples, JRW and EZ supervised research; JD and JRW conceived the project and prepared the manuscript; all authors commented on the manuscript.

---

**Supplementary Table 1:** Antibodies used in study.

| Antibody | Manufacturer | Application |
| --- | --- | --- |
| **GSK-3 (Total)** | Life Technologies | Immunoblot (IB) |
| **GSK-3β** | Cell Signaling Technology 9315 | Immunohistochemistry (IHC) |
| **β-catenin** | BD Transduction (mouse) | IB, IHC |
| **β-catenin** | Cell Signaling Technology 9581 (Rabbit) | IHC |
| **Keratin 8/18** | Fitzgerald | IHC |
| **Keratin 14** | Covance | IHC, Immunofluorescence (IF) |
| **Keratin 6** | Covance | IHC |
| **α-Smooth Muscle Actin** | Novus Biologics | IHC |
| **E-cadherin** | BD Transduction | IHC |
| **Ki67** | Biocare Medical | IHC |
| **Cyclin D1** | Calbiochem | IB |
| **BrdU** | abcam | IHC |
| **Cre** | Covance | IHC, IB |
| **EGFR** | Cell Signaling Technology 2646 | IHC |
| **γ-catenin** | Sigma clone 15F11, P8087 | IB |

**Supplementary Table 2:**
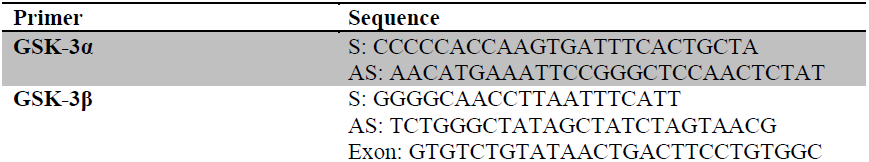
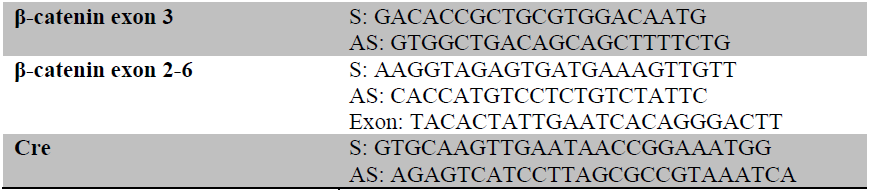
Genotyping PCR primers.

